# scMoE: single-cell mixture of experts for learning hierarchical, cell-type-specific, and interpretable representations from heterogeneous scRNA-seq data

**DOI:** 10.1101/2024.10.24.620111

**Authors:** Michael Huang, Yue Li

## Abstract

Advancements in single-cell transcriptomics methods have resulted in a wealth of single-cell RNA sequencing (scRNA-seq) data. Methods to learn cell representation from atlas-level scRNA-seq data across diverse tissues can shed light into cell functions implicated in diseases such as cancer. However, integrating large-scale and heterogeneous scRNA-seq data is challenging due to the disparity of cell-types and batch effects. We present single-cell Mixture of Expert (scMoE), a hierarchical mixture of experts single-cell topic model. Our key contributions are the cell-type specific experts, which explicitly aligns topics with cell-types, and the integration of hierarchical cell-type lineages and domain knowledge. scMoE is both transferable and highly interpretable. We benchmarked our scMoE’s performance on 9 single-cell RNA-seq datasets for clustering and 3 simulated spatial datasets for spatial deconvolution. We additionally show that our model, using single-cell references, yields meaningful biological results by deconvolving 3 cancer bulk RNA-seq datasets and 2 spatial transcriptomics datasets. scMoE is able to identify cell-types of survival importance, find cancer subtype specific deconvolutional patterns, and capture meaningful spatially distinct cell-type distributions.

## 1 Introduction

Transcriptomics is the study of Ribonucleic Acid (RNA) transcripts in organisms, allowing researchers to capture the expression of genes in order to understand their functions and study diseases [1, 2]. Bulk RNA sequencing (RNA-seq) is a technique that measures pooled gene expression levels within a tissue sample [3]. With advancements in technologies, bulk RNA-seq has become inexpensive, leading to large repositories of data widely available for researchers, such as The Cancer Genome Atlas (TCGA) [4]. A limitation of bulk RNA-seq is that it fails to capture cell-type resolution heterogeneity. Single-cell RNA sequencing (scRNA-seq) is a technique that allows researchers to profile single-cell resolution gene expression levels, enabling researchers to elucidate cell-type population differences previously masked by bulk sequencing [5]. scRNA-seq is a field that has grown significantly in past years, and has yielded results in Cancer [6], Alzheimer’s disease [7] and Covid-19 [8].

While single-cell transcriptomics provides high throughput, high resolution datasets, it is not without its drawbacks. The process of generating scRNA-seq requires the isolation of cells and tissue dissociation. Additionally, not all tissues can be easily processed for scRNA-seq [9]. Spatial transcriptomics (ST) is the technique whereby transcriptomics is performed on intact tissues. In addition to addressing dissociation difficulties, ST provides spatially resolved data, preserving positional information that would otherwise be lost, such as cell-cell signalling. Although ST is still limited in terms of gene-coverage and throughput compared to scRNA-seq, it has proven to be a rapidly evolving technology. While ST technologies such as MERFISH [10] and seqFISH+ [11] can achieve single-cell resolution, they only profile about one thousand genes. Commercial platform Visium can achieve transcriptome-wide profiling but are not single-cell resolution, resulting in spot-level transcriptome measurement where each spot typically contains 5-10 cells. This calls for deconvolution methods that can accurately dissolve the cell mixture in each spot into cell-type proportions.

As larger scRNA-seq datasets have become available, tools for integrative analysis have been developed to cluster cells for the identification of novel cell-types, correct for experimental batch effects, and transfer information to new datasets. Seurat [12] uses canonical correlation analysis to project cells onto an integrated embedding space, in order to maximally align cells across datasets. Harmony [13] uses principal components analysis in order to project cells onto a lower dimension, whereby it then alternates between clustering and batch correction. Recently, deep learning based approaches have proliferated greatly. scVI [14] is an unsupervised variational inference-based approach implemented using a variational autoencoder (VAE). It has inspired several extensions, such as LDVAE [15] and scETM [16], sporting a multilayer perceptron (MLP) encoder and an interpretable linear decoder, and scANVI [17], a semi-supervised extension that incorporates cell-type labels into the inference process.

In this study, we present single-cell Mixture of Experts (scMoE), a highly interpretable, flexible method for single-cell modeling with applications in clustering, deconvolution, and gene set enrichment analysis. We benchmarked scMoE on scRNA-seq data for clustering and on simulated bulk ST data for spatial deconvolution, against existing single-cell modeling and spatial deconvolution methods respectively. We then demonstrate deconvolution applications using single-cell references applied to various bulk RNA-seq cancer datasets. When applied to deconvolving bulk pancreatic cancer, scMoE successfully distinguishes subtypes by clustering on predicted cell-type proportions. When applied to deconvolving bulk lung cancer and bulk breast cancer, scMoE successfully identifies cell-type proportions of patient survival importance. Furthermore, on bulk breast cancer data we show scMoE can perform cell-type specific differential expression analysis between cancer subtypes and identify relevant biological pathways. Moreover, we show two applications of spatial deconvolution. When deconvolving a ST dorsolateral prefrontal cortex dataset, we show that scMoE is able to predict distinct layer-wise spatial cell-type distributions. We deconvolve a ST breast cancer dataset, and show that the predicted spot cell-type patterns are conserved across replicates. Furthermore, the deconvolved proportions capture differences between non-malignant spots and two types of malignant spots.

## 2 Methods

### 2.1 scMoE model description

scMoE consists of two extensions to the scETM model proposed by Zhao et al. [16]. The first is an extension to scETM using a traditional Mixture of Experts (MoE) approach proposed by Jacobs et al. [18], whereby pre-trained cell-type specific experts are modulated by a gating neural network. Next is a further extension that exploits the knowledge of hierarchical cell lineages, and the domain knowledge of cell-type lineage tree structures.

The scMoE model consists of *K* cell-type specific experts, each being its own scETM unit with an encoder and linear decoder. Additionally, the model contains *G* gating networks, corresponding to the specific number of hierarchical branches used for the model. scMoE generates a unified latent embedding by combining the expert encoders and modulating their values using its gating networks. The model interpretability is maintained by concatenating each cell-type expert’s linear decoder. This formulation results in topics being explicitly assigned specific cell-types a priori. The model training process is as follows: we first split the single-cell reference data into cell-type specific datasets, and pre-train localized experts on specific cell-types. We then fix the expert parameters, and fine-tune the gating networks on a target dataset to integrate the experts. An overall model diagram is illustrated in Fig. 1a. The details are described as follows.

**Figure 1:**
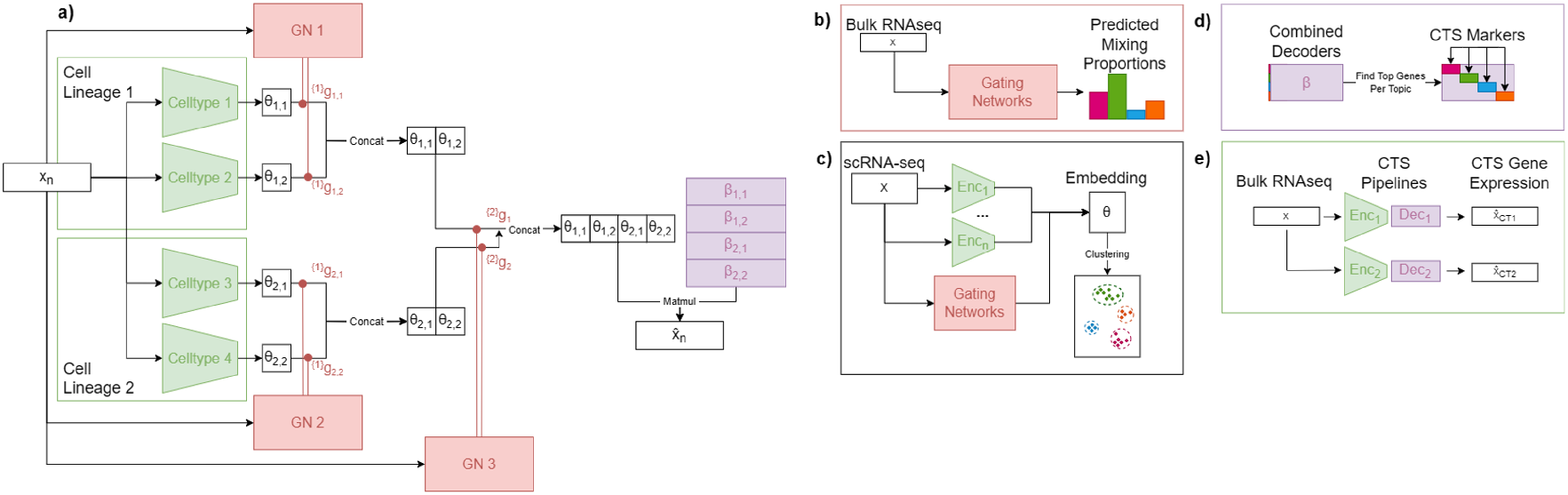
scMoE model overview. (a) Hierarchical scMoE architecture. scMoE consists of pre-trained cell-type specific (CTS) experts (illustrated as green encoders and purple linear decoders) and a flexible set of hierarchical gating networks (illustrated in red). (b) Gating network weights can be used for bulk RNA-seq deconvolution, output weights are equivalent to predicted proportions. (c) Latent embedding *θ* can be used for clustering and visualization. (d) Combined linear decoders provide interpretability, top genes per topic correspond to CTS markers. (e) CTS encoder/decoder pairs can be used to generate cell-type specific gene expression profiles from bulk RNA-seq data.

#### 2.1.1 Single-cell embedded topic model and variational autoencoder

Each expert unit consists of a single-cell embedded topic model (scETM). scETM is an unsupervised generative topic model proposed by Zhao et al. [16]. The model consists of an encoder and linear decoder. Assume a dataset of cells *X* = {*x*_1_, …, *x*_*D*_} ∈ ℝ^*D×G*^ with *D* cells and *G* genes. The linear decoder is parameterized by *α* ∈ ℝ^*T*×*L*^, *ρ* ∈ ℝ^*L*×*G*^, where *T* is the number of latent topics, *L* is the embedding size, and *G* is the number of genes. The encoder is a feed-forward neural network, FFN(*x*_*d*_; *W*_*θ*_) = *µ*_*d*_,Σ_*d*_. The reparameterization trick is then used to get the unnormalized latent embedding δ_*d*_. The normalized embedding *θ*_*d*_ is then computed via Softmax:

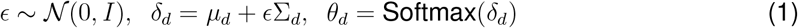

where *θ*_*d*,*t*_ = exp(δ_*d*,*t*_)*/*Σ_*t*_exp(δ_*d*,*t*_) for *t* ∈ {1, …, *T* }.

For cell *d*, the reconstructed normalized gene expression 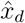 is obtained using the following:

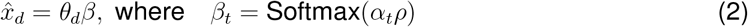

where *β*_*t*,*g*_ = exp(*α*_*t*_*ρ*_*g*_)*/*Σ_*g*_exp(*α*_*t*_*ρ*_*g*_) for *g* ∈ {1, …, *G*}

In order to learn the parameters *W*_*θ*_, *α, ρ*, we optimize the Evidence Lower Bound (ELBO), where the first term is the reconstruction loss, and the second term is the Kullback-Leibler divergence:

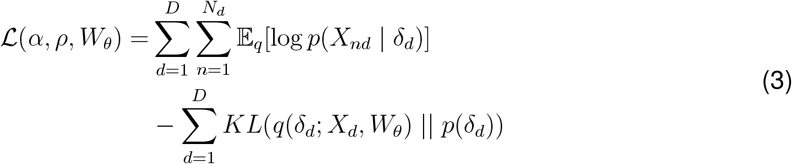

A limitation of scETM is that it requires post-hoc analysis of topics in order to investigate cell-type specific gene signatures.

#### 2.1.2 Mixture of experts formulation

To address the above challenge, we present scMoE. This section describes a basic scMoE model with no hierarchical information and the next section describes the hierarchical version. In scMoE, each expert is an scETM unit trained on a known cell-type, thus explicitly partitioning the problem space and enabling us to directly assign cell-types to topics. Specifically, consider a dataset of single cells *X* = {*x*_1_, …, *x*_*D*_}, which we then partition into *K* unique cell-types, where *X*_*k*_ is the dataset partition only containing cells of cell-type *k*. We then train *K* different scETM experts, one for each cell-type, each with a latent dimension *L*. We then combine the scETM units together, using a gating network GN(*x*) = *g* ∈ ℝ^*K*^. We can formalize the reconstruction process as follows:

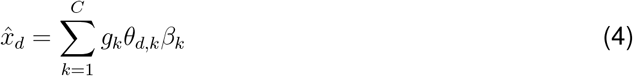

where *θ*_*k*_ is the latent representation generated by the k-th encoder, *β*_*k*_ is the k-th linear decoder, and *g*_*k*_ is the k-th index output of the gating network. During the integration of cells across diverse cell-types, we freeze the cell-type-specific expert parameters *W*_*θ*_, α, *ρ* and fine-tune only the gating network parameters *W*_*g*_.

#### 2.1.3 Hierarchical mixture of experts formulation

Hierarchical MoEs were proposed as a direct extension to MoEs by Jordan et al. [19]. Likewise, we propose an extension to scMoE to include hierarchical information. The above scMoE can be considered as a special case with a single hierarchical level. The hierarchical scMoE is a tree, where each leaf is a localized scETM expert, and each internal node determines some cell-type grouping. An internal node *N* contains a gating network, which consists of: a feed-forward neural network with a fixed output size |*g*| = |Children(*N*)| (i.e. the number of children) and a Softmax layer. In our setting, we can consider the outputs of the gating network at each node to be the probabilities assigned to all of its children, and thus the final weight assigned to a leaf is the cumulative product of all weights tracing the path from the root to the leaf. Let {*j*|*k*} be the set of nodes and their associated gating networks *N*_*j*_ between an expert *k* and the root of the tree. The reconstructed gene expression for cell *d* is as follows:

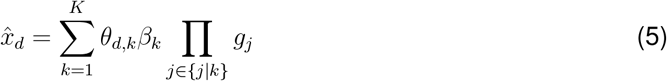

### 2.2 Performance Evaluation

We compared scMoE to three state of the art, deep learning, variational inference-based, single-cell modeling methods: scVI [14], scANVI [17], and LDVAE [15]. We performed 5-fold cross validation, on 9 scRNA-seq datasets, all datasets are detailed in Table S1. We evaluated the model clustering performance using three standard metrics. Specifically, we used the Adjusted Rand Index (ARI) and Adjusted Mutual Information score (AMI) in order to evaluate the pairwise labeling agreement between predicted cell-type labels and ground truth labels. The predicted cell-type labels are generated by computing the Louvain [20] clusters on the latent embeddings. We additionally computed the Average Silhouette Width (ASW), using the ground-truth cell-type labels and latent embeddings, without the use of any additional clustering algorithm.

We additionally compared scMoE to several published spatial deconvolution methods including Tangram [21] and scProjection [22], which are deep-learning based models. We also compared scMoE to RCTD [23], a probabilistic deconvolution method. In order to evaluate performance on spatial deconvolution, we simulated bulk spatial data using MERFISH single-cell resolution spatial data. We evaluated model performance on the simulated spatial data using the Spearman’s Correlation, Pearson’s Correlation, and Root Mean Squared Error (RMSE) between the predicted bulk mixing proportions and ground-truth simulated proportions.

### 2.3 Pathway enrichment analysis

Gene set enrichment analysis (GSEA) was performed using the gseapy [24] python library, using two pathway sets: the MSigDB HALLMARK 2020 [25,26] and the 2023 GO Biological Processes [27, 28].

### 2.4 Cell-type specific differential expression analysis

In order to leverage the cell-type specific (CTS) experts for differential expression, we can consider the predicted gene expression of a CTS expert for a bulk RNA-seq sample to be the CTS gene expression. For cell-type *k*, the predicted CTS gene expression is 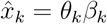. We can then perform a two-sided T-test for each CTS predicted gene expression for two populations.

### 2.5 Survival analysis

For bulk RNA-seq TCGA datasets, expert associated gating-weight values were used as cell-type proportion predictions. Survival analysis is performed using the sksurv python package [29]. Samples were split into binary categories by predicted cell-type proportion (cutoffs from 0.1 to 0.5, in increments of 0.1). We performed k-sample log-rank tests using the proportion cutoffs. The resulting p-values were adjusted by Bonferonni correction for multiple testing. For the cell-type proportion with *q <* 0.05, we examined their Kaplan-Meier curves.

### 2.6 Experimental setup

#### 2.6.1 Data preprocessing

For gene pre-processing, we removed all ribosomal, mitochondrial, and non-coding genes from the single-cell reference. We additionally removed cells with less than 200 counts, and genes that appeared in fewer than 3 cells. When using single-cell as reference for a bulk RNA-seq dataset, only overlapping genes were used.

#### 2.6.2 Quantitative benchmark

For single-cell quantitative evaluation, we performed 5-fold cross-validation. Each dataset was split into 5 partitions, where for each split, 4 partitions are used as training data, and 1 partition is withheld as test data. During training, the data were further split into 90-10 training and validation, where the validation data were used for early stopping to prevent overfitting. The training data were then partitioned into cell-type specific subsets, and the experts were individually trained on their respective subsets. The expert weights were then frozen, and the gating weights were fine-tuned on the merged training data. Finally the latent embeddings for the heldout partition were computed, and clustering metrics were recorded.

#### 2.6.3 Bulk deconvolution

When applying scMoE to deconvolve bulk RNA-seq data, we used single-cells as reference for expert pre-training. We then fixed the experts and minimized reconstruction loss on the bulk RNA-seq samples. We used this training setup for the bulk RNA-seq TCGA cancer data, the simulated spatial bulk data, and the real ST datasets.

#### 2.6.4 Simulation of single-cell spatial transcriptomes

In order to simulate spatial pseudobulk data, we partitioned the MERFISH single-cell spatial data into 100×100 pixel bins for the Mouse Medial Preoptic Area and Mouse Dorsolateral Prefrontal Cortex datasets, and 1000×1000 pixel bins for the Mouse Kidney dataset. The counts for each bin were then summed, and the proportions of single-cells composing those bins were used as the ground truth mixing proportions. The predicted mixing proportions are the gating weights obtained from the fine-tuned scMoE model on the simulated bulk data. The data simulation process is illustrated in Fig. 2.

**Figure 2:**
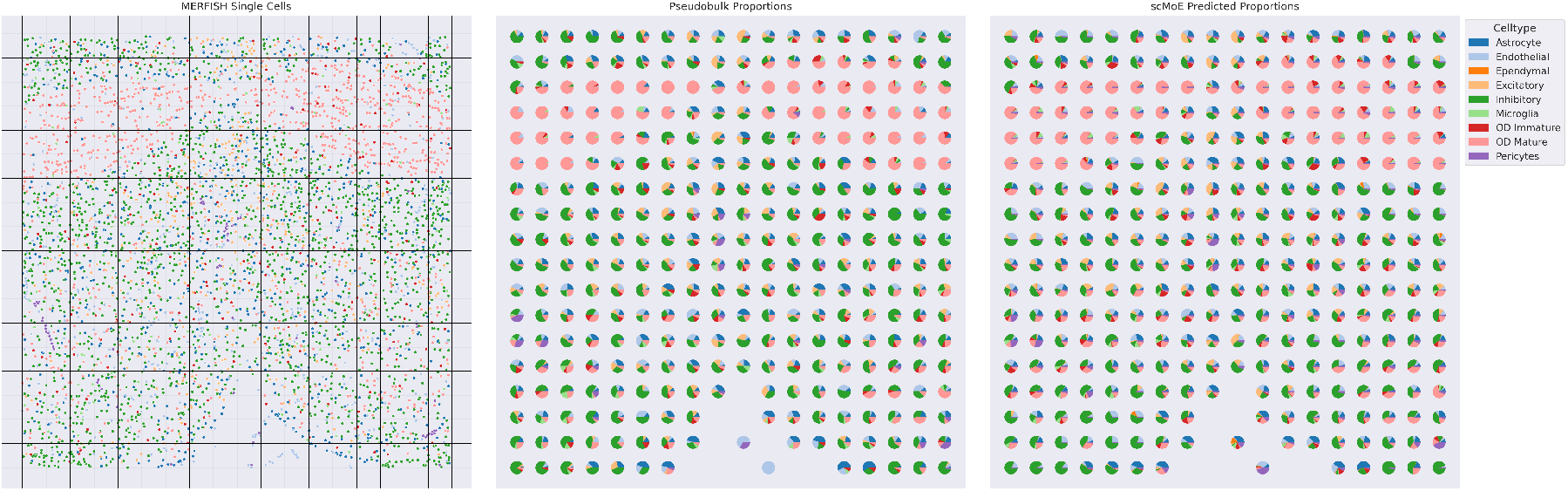
Spatial pseudobulk simulation process (MPOA shown). Spatial slide is segmented into 100×100 pixel bins. Simulated spots with less than 5 cells are dropped. Spot gene expression profiles are generated by summing the MERFISH single-cell profiles within each bin, cell-type proportions are recorded as the ground truth. Left panel shows MERFISH single-cell labels partitioned into bins. Middle panel shows ground truth mixing proportions of simulated spots. Right panel shows scMoE predicted proportions.

### 2.7 Model Implementation

Each scETM encoder and each internal gating network had two fully connected feed-forward layers (128, 128), BatchNorm, Dropout (dropout probability = 0.1), and ReLU nonlinearity layers. The gating networks contain an additional feed-forward layer and Softmax function. Each cell-type specific expert was given a latent dimension of 5. The topic and gene embedding dimensions were set to 400. The Negative ELBO was minimized using the Adam optimizer [30] as implemented in PyTorch. The learning rate for cell-type specific model training was 1e-3, while the learning rate for gating-network fine-tuning was 1e-4. Detailed cell-type hierarchies used for each dataset is described in detail in Appendix Fig. S3-S4.

## 3 Results

### 3.1 Single-cell clustering benchmarking

We benchmarked scMoE against three existing single-cell modeling methods – scVI [14], scANVI [17], and LDVAE [15]. We also compared an ablated scMoE model, which has all experts in a single layer, with a single gating network, which we label scMoE-H. We compare these methods using 5-fold cross validation, on 9 published single-cell datasets: Alzheimer’s Disease dataset (AD) [7], Adult Human Heart (AHH) [31], COVID-19 immune cells (COVID) [32], Human Breast Cell Atlas (HBCA) [33], Human Cell Landscape (HCL) [34], Hematopoietic Immune Cell Atlas (HICA) [35], Human Lung Cell Atlas (HLCA) [36], Major Depressive Disorder dataset (MDD) [37], and the Tabula Sapiens dataset (TS) [38]. For TS and HCL, only cell-types with 1000 cells after quality control were used, for the other 7 datasets, all cell-types in the datasets were used. Both scMoE and scMoE-H outperform existing methods on 5 of 9 datasets in terms of both ARI and AMI, and is competitive with existing methods on the remaining 4 datasets (Fig. 3). In terms of ASW, both scMoE and scMoE-H significantly outperforms existing methods.

**Figure 3:**
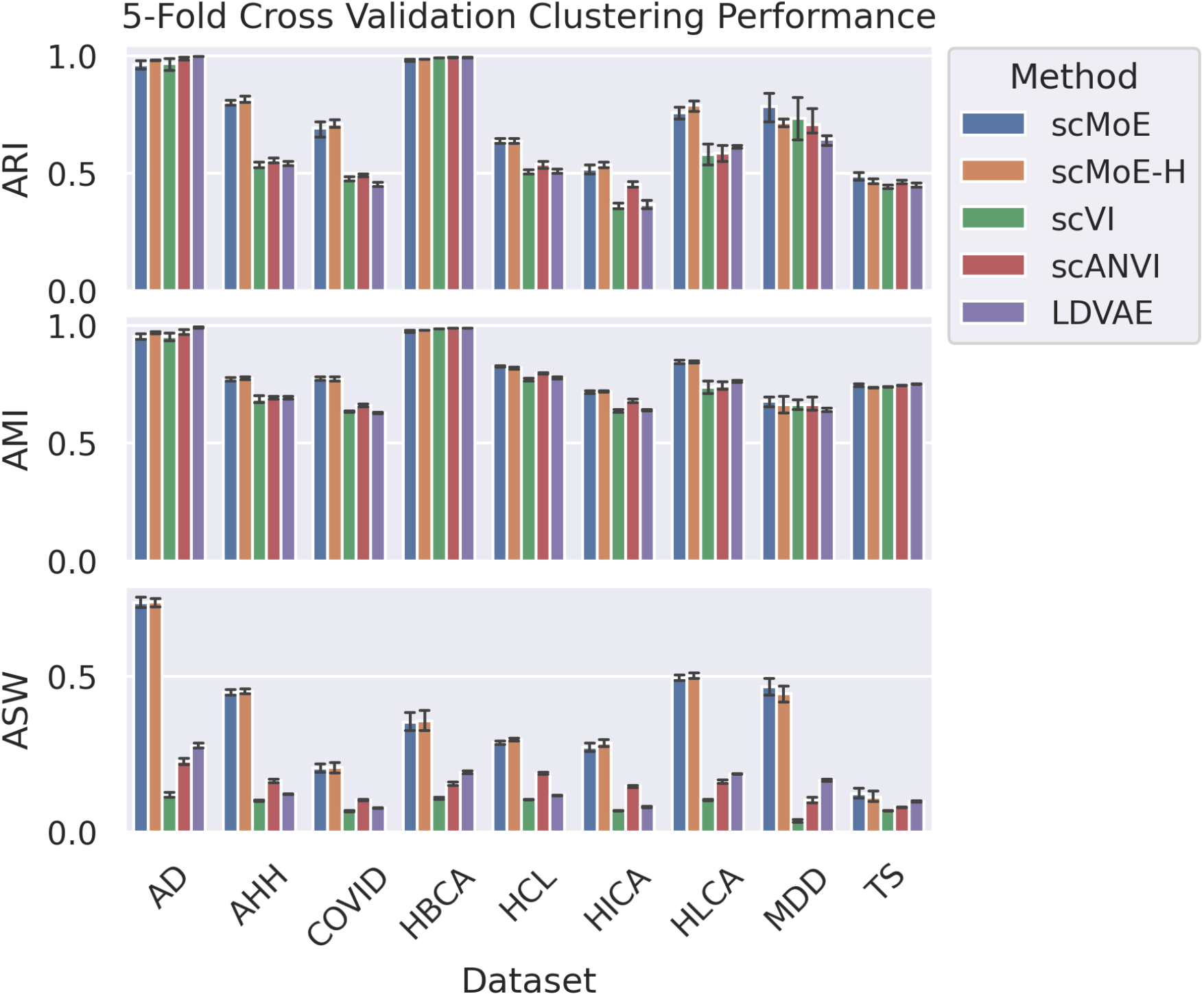
5-fold cross validation quantitative single-cell clustering benchmarking results. Adjusted Rand Index (ARI), Adjusted Mutual Information (AMI), and Average Silhouette Width (ASW) were computed using each methods latent embedding representations.

To further verify the quantitative clustering metrics, we visualized the latent embeddings and the resultant hierarchical clustering using Seaborn [39] for the AD dataset (Appendix Fig. S1). These heatmaps highlight scMoE’s improved interpretability compared to existing methods. There is clear cell-type specific alignment of topics. Additionally the hierarchical clustering shows clearer cell-type clustering compared to scVI, scANVI, and LDVAE.

### 3.2 Model Interpretability

In order to highlight the interpretability of scMoE’s decoder, we sorted the combined linear decoder by top 5 genes per topic and compared the top genes against known cell-type markers in CellMarkerDB [40]. We illustrate the results of scMoE trained on the AD dataset for the first 4 cell-types in Fig. 4a. There is a clear enrichment for corresponding cell-type marker genes and their known cell-type expert topics.

**Figure 4:**
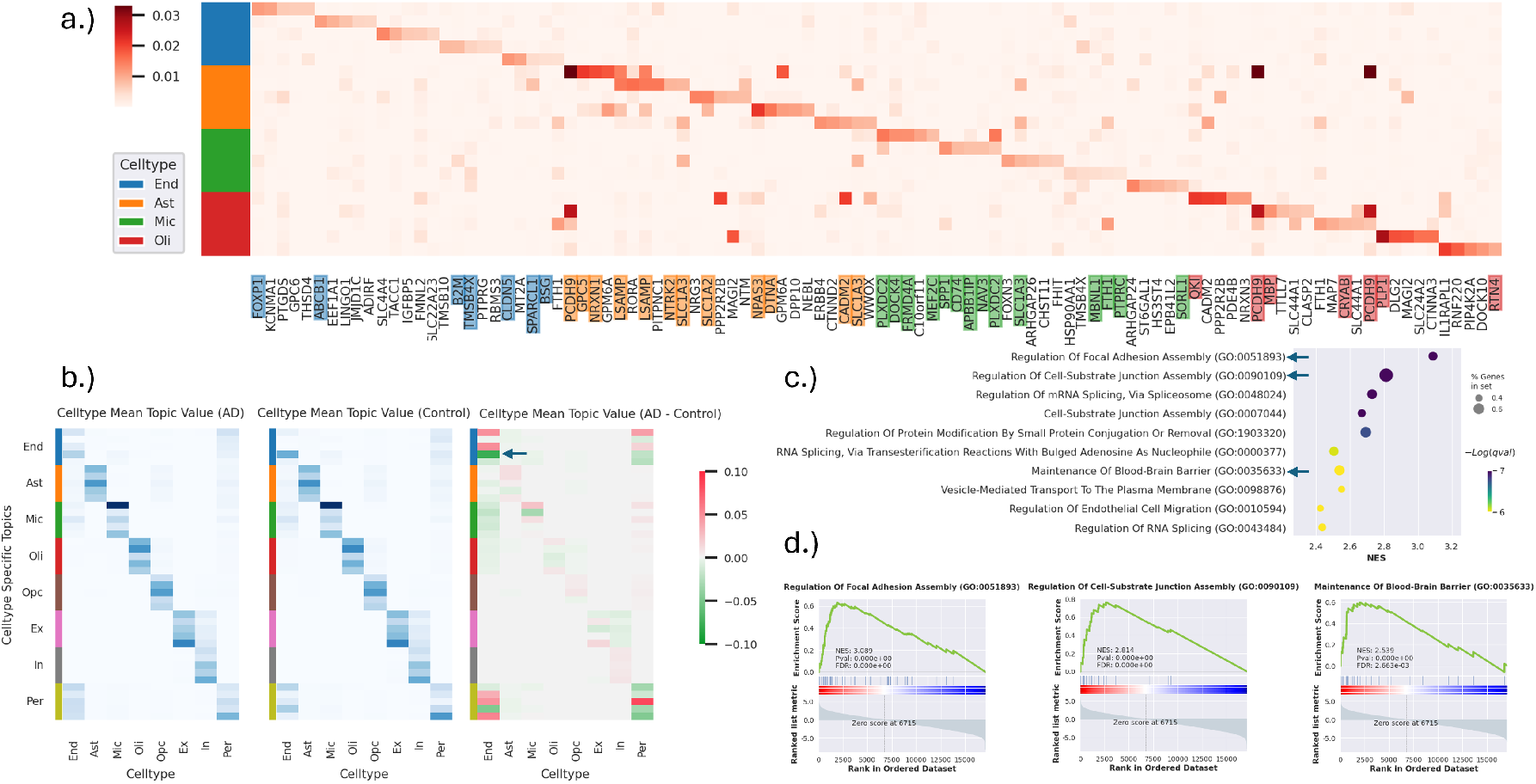
Alzheimer’s disease model decoder interpretability. (a) Heatmap for the linear decoder *β*, showing top 5 genes per topic, genes matching CellMarkerDB cell-type marker genes are colored accordingly. Topics 1-20, corresponding to endothelial, astrocyte, microglial, and oligodendrocytes are illustrated. (b) Mean cell-type topic density for withheld cells (Cross Validation Fold 1), comparing AD cells, control cells, and the difference: AD - Control. Colors correspond to cell-type specific topics. (c) Top 10 enriched pathways for topic 4, an endothelial cell topic. (d) Select relevant gene set enrichment leading edge curves, elevated in control relative to AD.

We then investigated the scMoE topics for cell-type specific biologically relevant pathways. In Fig. 4b. we compared the cell-type specific topic representations for both AD and Control cells. We highlighted Topic 4, which is highly represented in endothelial cells from control subjects compared to those from AD (Fig. 4b,c). We observe pathways that are highly relevant to endothelial cells and their function in the brain. We also highlighted the “Maintenance of Blood-Brain Barrier” pathway, which does not appear in the top 30 pathways for Topics 1 and 3, which are conversely highly represented in AD compared to control.

### 3.3 Deconvolving pancreatic bulk RNA-seq using Human Cell Atlas as reference identifies tumor subtype of PAAD

We trained an scMoE model on the HCL single-cell atlas as reference for deconvolving TCGA pancreatic adenocarcinoma (PAAD) bulk RNA-seq data. Fig. 5 displays the heatmap clustering based on the predicted proportions by the gating network. We were able to clearly identify a distinct subset of samples: the neuroendocrine subtype, which has a unique and distinctly high predicted proportion of astrocytes.

**Figure 5:**
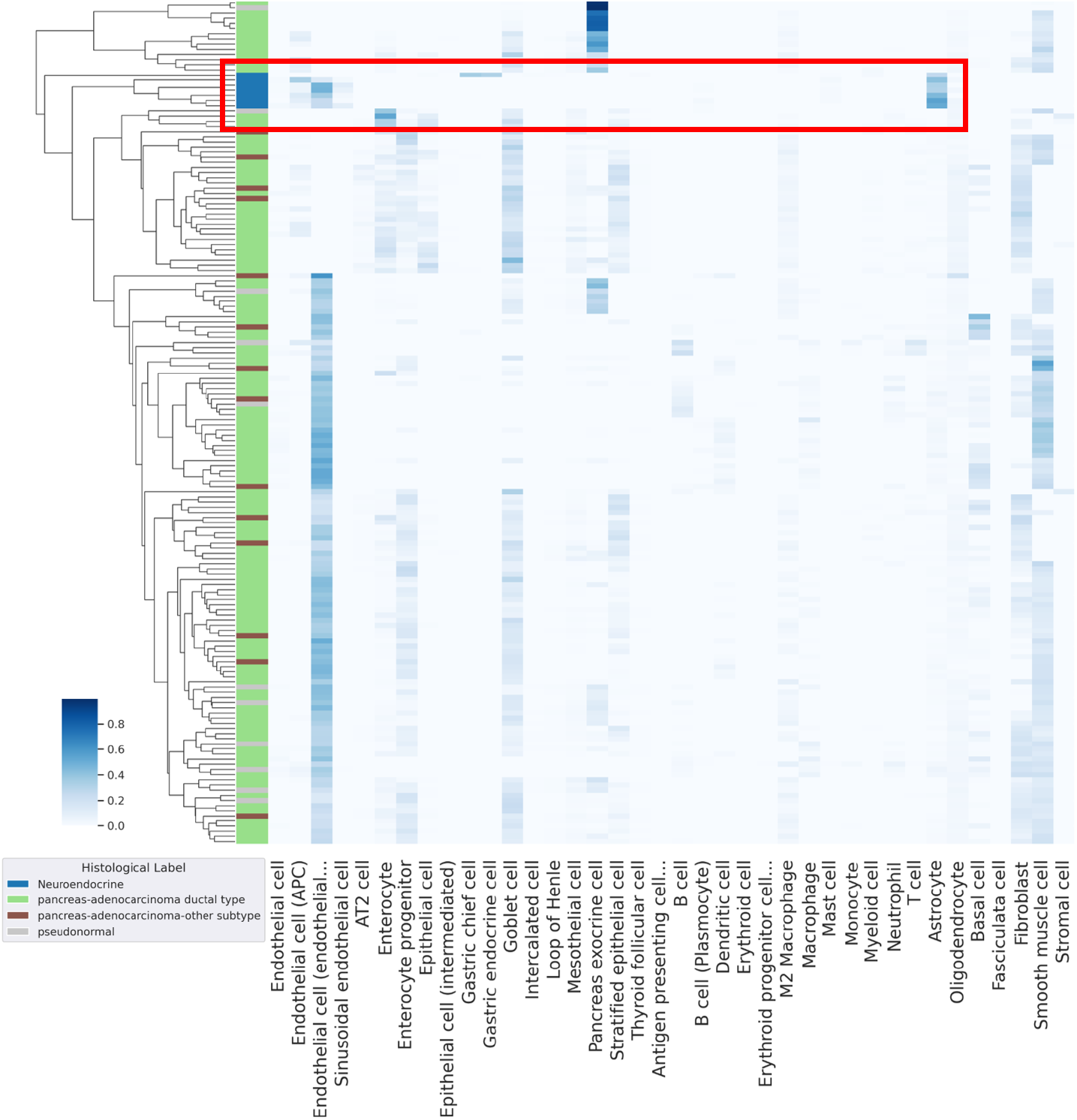
Bulk pancreatic cancer deconvolution using HCL as reference. Row color labels indicate histological subtype. Heatmap intensity shows scMoE predicted cell-type proportions. Red box highlights the neuroendocrine subtype.

### 3.4 Predicted cell-type proportions as survival indicators in lung cancer samples

We trained an scMoE model with the HLCA as reference and applied it to the TCGA lung adenocarcinoma (LUAD) bulk RNA-seq dataset. We performed survival analysis by separating tumor samples based on gating weight predicted threshold proportions, and found two cell-types conferring significant Bonferroni corrected p-values (Fig. 6a): fibroblasts (p-value = 1.3717e-4, log-rank test), and basal cells (p-value = 1.6774e-04, log-rank test). In order to further support these findings, we performed paired t-tests using normal and tumor tissues sampled from the same individuals in the TCGA-LUAD [41] dataset. We projected the predicted bulk RNA-seq proportions using t-SNE [42] and visualized the differences in predicted proportions between the healthy and tumor tissues. The Fibroblasts (p-value = 3.251e-9, paired t-test) and Basal cells (p-value = 3.561e-7, paired t-test) both confer significant p-values, which is clearly reflected in the predicted proportions (Fig. 6b).

**Figure 6:**
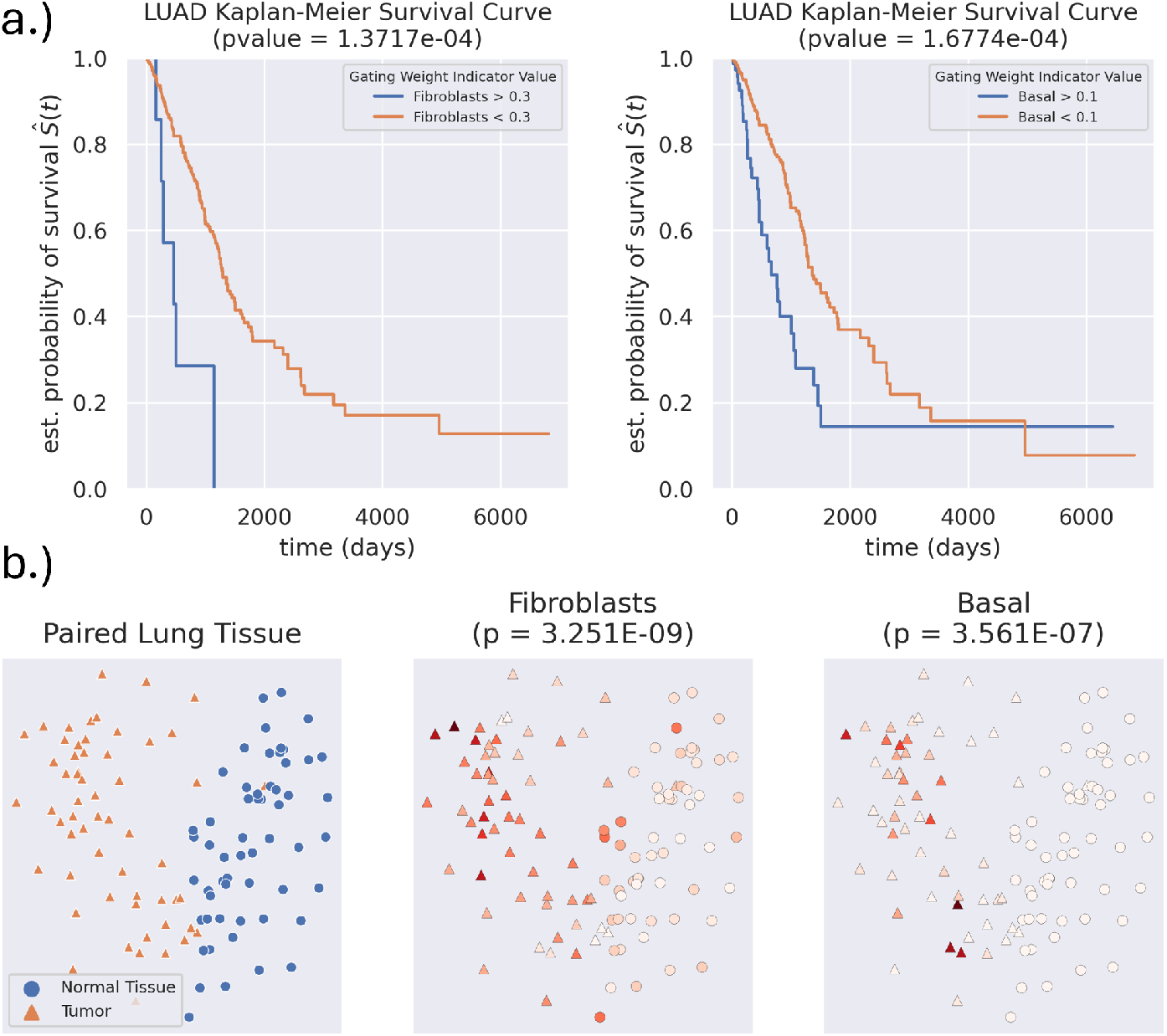
Bulk lung adenocarcinoma deconvolution using HLCA as reference. (a) Kaplan-Meier survival curves computed using lung tumor samples, fibroblasts and basal cells have significant p-values after Bonferroni correction. (b) Patient paired normal and tumor tissue t-SNE plot. First panel shows sample status, remaining panels show predicted relative cell-type proportion.

### 3.5 Cell-type-specific differential gene expression analysis identifies pathways indicative of breast cancer subtypes

We trained an scMoE model with the HBCA as reference and applied it to the TCGA breast cancer bulk RNA-seq dataset. We observe a strong subtype clustering pattern when utilizing the gating network predicted proportions, where the basal subtype has a distinctly higher predicted proportion of myoepithelial cells (Fig. 7a). We additionally performed Kaplan-Meier survival analysis and found 4 cell-types to be statistically significant after Bonferroni correction (blood vessel endothelial cells, and the fibroblast, myoepithelial, and luminal epithelial cells of the mammary gland, Appendix Fig. S2).

**Figure 7:**
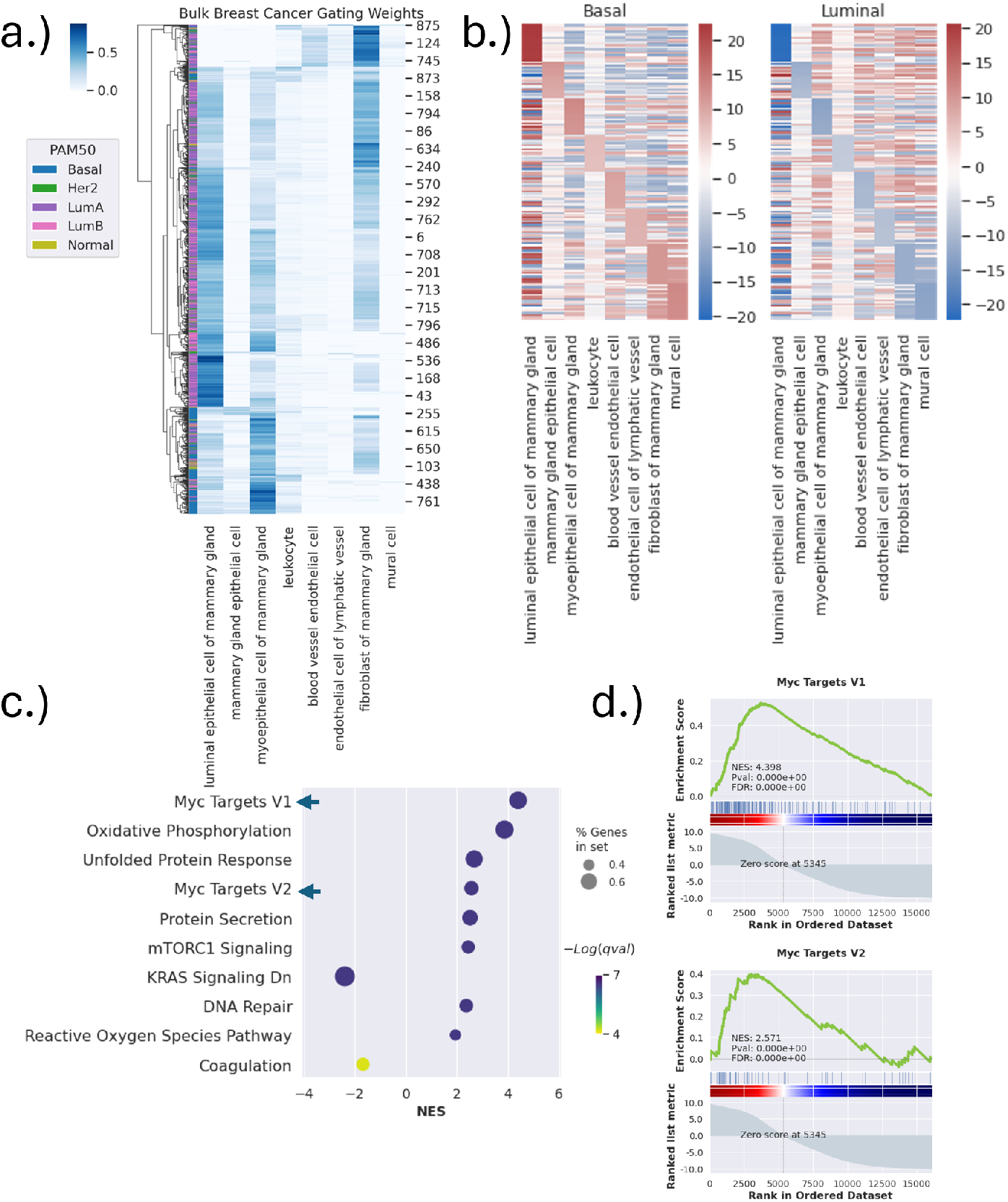
Bulk breast cancer deconvolution using HBCA as reference. (a) Heatmap containing the predicted cell-type proportions of bulk cancer data, row clustering color index indicates breast cancer subtype. (b) Heatmap comparing CTS and DE gene expression between Basal and Luminal subtypes. Top 20 DE genes in Basal subtype relative to Luminal subtypes for each of the 8 cell-type experts are shown. (c) Top 10 differential MSigDB HALLMARK 2020 pathways by p-value for the BVEC cell-type, with Myc Targets V1/V2 highlighted. (d) Gene set enrichment leading edge curves for Myc Targets V1/V2.

We then performed CTS differential expression (DE) analysis using the imputed CTS gene expression by scMoE, comparing basal and luminal breast cancer subtypes. Specifically, each cell-type expert in scMoE was used to predict CTS gene expression given the same bulk RNA-seq data as the input. We performed independent t-tests for each cell-type and ranked genes by their t-statistic scores. The top 20 DE genes per cell-type enriched in Basal compared to Luminal subtypes are visualized in Fig. 7b. Pathway enrichment analysis of the “Blood Vessel Endothelial Cells” using MSigDB HALLMARK 2020 [25] reveals both Myc Targets V1 and Myc Targets V2 as enriched in Basal relative to the Luminal breast cancer samples (Fig. 7c,d).

### 3.6 Spatial deconvolution benchmarking

We compared scMoE and scMoE-H (i.e., without hierarchy) against 3 existing spatial deconvolution methods: scProjection [22], RCTD [23], and Tangram [21]. We benchmarked on simulated data based on 3 MERFISH single-cell resolution ST datasets: Mouse Medial Preoptic Area (MPOA) [43], Mouse Brain Aging Spatial Atlas (MBASA) [44], and Mouse Spatial Kidney (MSK) [45]. The MPOA, MBASA, and MSK used MERFISH (59541 cells, 156 genes) [43], snRNA-seq (79328 cells, 374 overlapping genes) [44], and scRNA-seq data (20441 cells, 291 overlapping genes) [38] as reference respectively. Data simulation process and scMoE predicted proportions for MPOA are shown in Fig. 2. scMoE performs equivalently, or better than all other methods in terms of Pearson R, and is either comparable or the second best in terms of Spearman R (Fig. 8). Compared against the ablation model, the hierarchical scMoE greatly outperforms scMoE-H in all metrics for the MPOA and MSK dataset, and has equivalent performance on the MBASA dataset.

**Figure 8:**
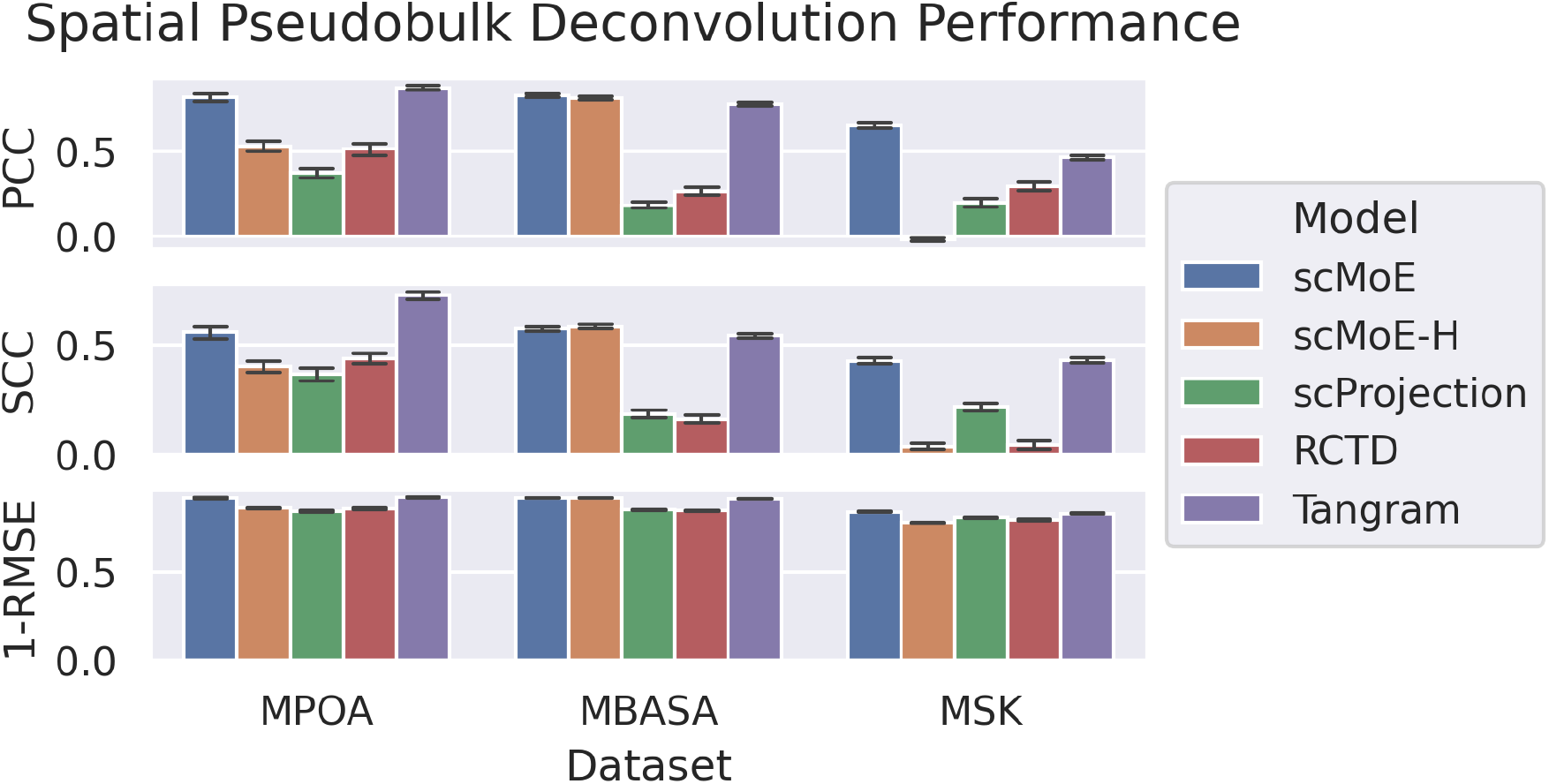
Spatial simulated pseudobulk deconvolution benchmarking. Models were trained on single-cell references and predicted proportions of simulated pseudobulk tissues. Known ground-truth proportions were used to compute Pearson’s Correlation Coefficient, Spearman’s Correlation Coefficient, and RMSE (In panel 3, 1-RMSE is illustrated to maintain directional coherence with SCC and PCC).

### 3.7 Spatial deconvolution reveals clear brain layer differentiation

We next performed spatial deconvolution of a real ST human dorsolateral pre-frontal cortex data generated by the Visium platform [46]. We trained scMoE using a single-cell prefrontal cortex dataset (AD) [7] as reference. The model gating networks were fine-tuned on all spots from a single slide. We visualized the predicted cell-type proportions in Fig. 9. We found clear localization of endothelial cells to the L1 and WM layers, astrocytes in the L1 layer, oligodendrocytes in the WM layer, and inhibitory and excitatory neurons in the L2-L6 layers.

**Figure 9:**
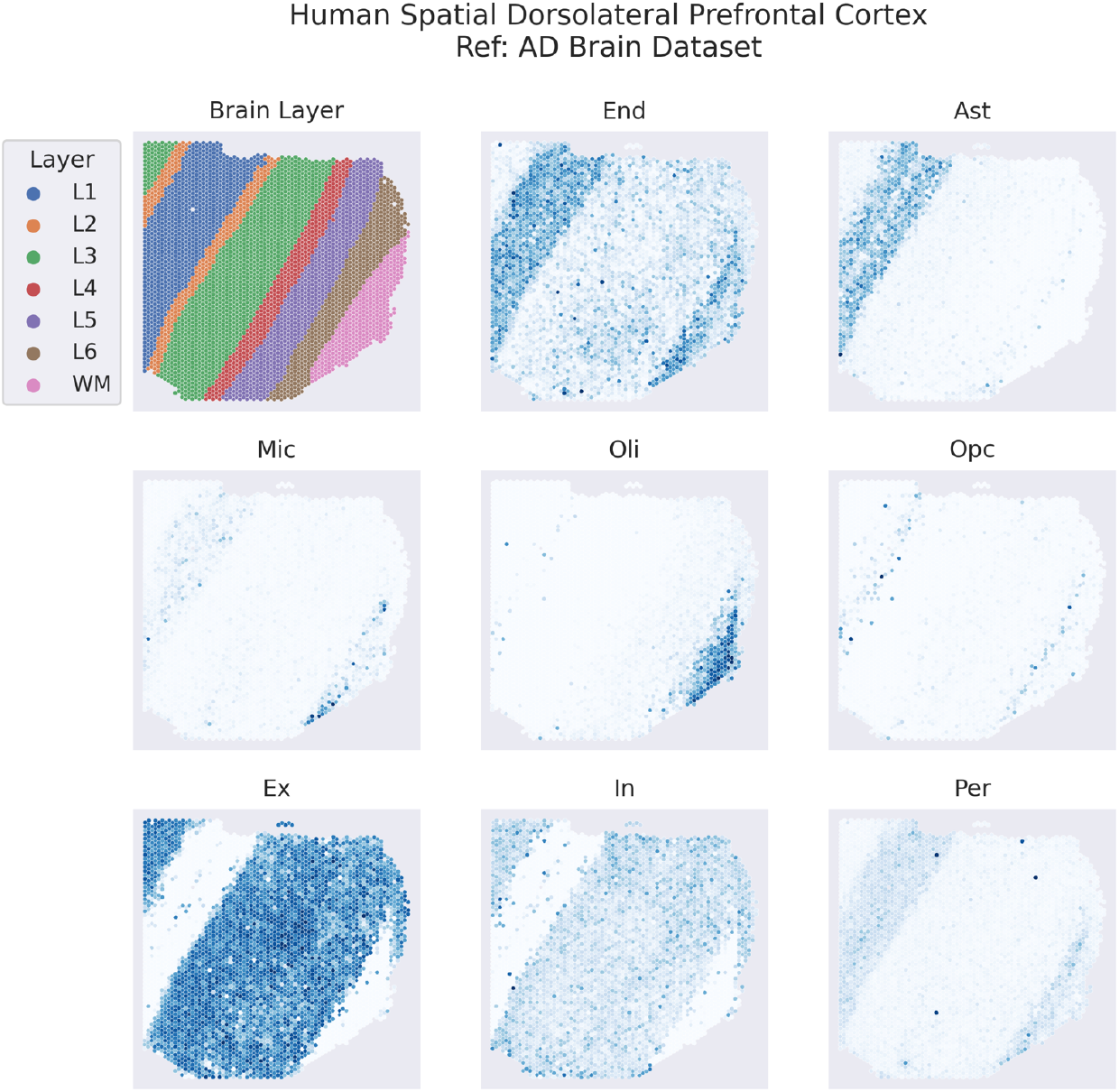
Predicted cell-type proportions for spot-resolution spatial dorsolateral prefrontal cortex data, using AD as reference. Top left panel: Brain Layer annotations. Remaining panels: Celltype specific scMoE predicted proportion for the spot-resolution spatial data.

### 3.8 Spatial deconvolution of breast cancer data yields distinct cell-type proportion pattern

We performed spatial deconvolution of ST breast cancer data, using the HBCA as a single-cell reference. The spatial breast cancer data consists of 4 slices of breast tissue. The data consists of 1031 spots, of which 194 were previously manually annotated by experts [47], identifying non-malignant, invasive ductal carcinoma (IDC), and ductal carcinoma in situ (DCIS) spots in the breast tissue. We observe distinct patterns in the deconvolution results generated by scMoE seen in Fig. 10a, which are conserved across slices.

**Figure 10:**
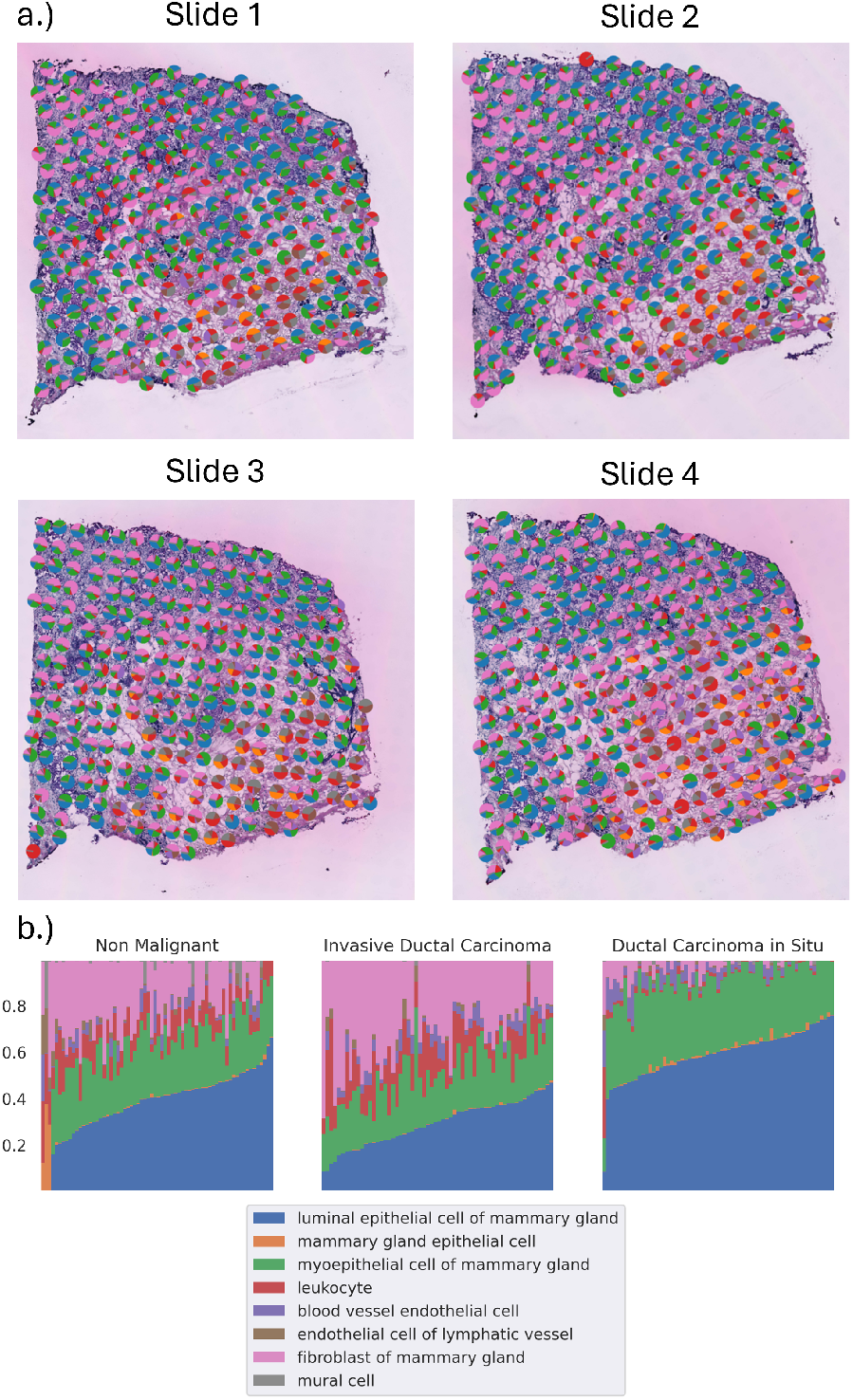
Spatial breast cancer deconvolution using HBCA as reference. (a) Spot-wise predicted cell-type proportions. Each spot is a pie-chart representing the scMoE predicted cell-type proportions. (b) Stacked bar charts representing the predicted proportions for spots that have expert annotations available, separated by classifications: non-malignant, IDC, and DCIS.

Using the expert annotations of the non-malignant, IDC, and DCIS spots, we were able to directly compare and visualize differences between the predicted cell-type proportions of each spot, as seen in Fig. 10b. There is an increase in fibroblast proportions and a slight increase in predicted leukocytes in IDC relative to non-malignant tissue. There is a larger difference when comparing the IDC and DCIS spots, in IDC there is an increase in fibroblasts, whereas DCIS spots have almost no predicted fibroblasts. The predicted proportion of luminal epithelial cells and myoepithelial cells are significantly increased in the DCIS spots, compared to both IDC and non-malignant spots.

## 4 Discussion and Conclusion

Single-cell transcriptomes provide rich information to dissect cell-type compositions from heterogeneous samples, associate marker genes, and identify cell-type-specific and differentially expressed genes indicative of cancer survival and cancer subtypes. Existing methods are limited either in their capacity to capture diverse cell-types or in their interpretability to distill meaningful biology. In this study, we developed a flexible mixture of experts single-cell topic model called scMoE, to infer interpretable, cell-type specific topic distributions and perform tasks such as cell-type clustering, cell-type specific differential expression analysis, and spatial deconvolution. We additionally incorporate hierarchical information, which utilizes further biological domain knowledge. To demonstrate scMoE, we conducted a comprehensive set of analyses using bulk and spatial transcriptome datasets. We found that incorporating hierarchical information improved quantitative on spatial deconvolution, but didn’t meaningfully change performance in terms of single-cell cell-type clustering (Fig. 3, 8).

We show that our model is highly interpretable, can easily identify cell-type marker genes (Fig. 4a), and identify biologically relevant pathways (Fig. 4c, 7d). Instead of post hoc analysis required by other interpretable models such as LDVAE to associate latent dimensions with cell-types, topics are immediately cell-type specific because each expert is pre-trained on unique cell-types. Through our analysis of the Alzheimer’s disease dataset, we are able to observe clear marker gene enrichment (Fig. 4). We are also able to identify cell-type specific pathways of relevance (Fig. 4bcd), and show that the blood-brain barrier (BBB) pathway is differentially reduced in Alzheimer’s endothelial cells relative to control cells. In particular the MSigDB HALLMARK pathway for maintenance of BBB is found in topic 4 which is differentially elevated in control relative to AD cells, corresponding to endothelial cells. Simultaneously the maintenance of BBB pathway is not in the top 30 pathways enriched in either topic 1 or 3, which are conversely elevated in AD relative to control. This is supported by several studies implicating BBB breakdown in Alzheimer’s patients [48, 49].

Through our analysis of the TCGA pancreatic cancer dataset, we show that our model’s interpretable outputs can cluster the Neuroendocrine subtype (Fig. 5), which was noted to have been previously misidentified [50]. We note that the Neuroendocrine cancer subtypes have a distinct, high predicted proportion of astrocytes, a neural cell-type present in the HCL reference data. This shows the utility of applying scMoE trained with large single-cell reference atlases. This would be particularly useful when a tissue specific reference may not be sufficient or available, whereby broad cell atlases or analogous references may be of interest.

Through our analysis of the TCGA lung cancer dataset, we show that our model’s deconvolution proportions can be used to identify cell-types of clinical relevance. Survival analysis found that elevated predicted proportions of fibroblasts and basal cells resulted in worse survival outcomes (Fig. 6a). These findings are further supported by the paired tumor and normal tissue samples (Fig. 6b), where scMoE deconvolution predicted elevated fibroblast and basal cell proportions in tumor samples compared to normal lung tissue. Both of these results are supported by studies regarding fibroblast [51] and basal cell [52] roles in lung cancer.

Through our analysis of the TCGA breast cancer dataset, we show that our model is able to separately cluster basal and luminal subtypes. We were also able to identify cell-types of survival relevance: increased proportions of fibroblasts and blood vessel endothelial cells resulted in worse survival outcomes, both cell-types highlighted in the literature [53, 54]. Enabled by the cell-type specific experts, we were additionally able to perform cell-type specific differential gene expression analysis and compare basal against luminal subtypes, and found meaningful pathway differences between the two types. In particular, BVEC pathways elevated in Basal subtype samples compared to Luminal subtypes include the Myc Targets V1 and Myc Targets V2 pathways. MYC has been found to be highly expressed in Basal subtypes but low in Luminal subtypes [55]. These results, taken together with our findings with the Pancreatic Neuroendocrine subtype, show that scMoE can be used to investigate disease subtypes.

We additionally show the utility of our model in a spatial transcriptomics setting. On bulk spatial dorsolateral prefrontal cortex data, we show that scMoE predicted cell-type proportions clearly identify layer specific cell-type distributions (Fig. 9). In particular, L1 and White Matter are distinctly different from the intervening layers. These findings are in concordance with known layer-specific cell-type distributions [11]. On ST breast cancer data, scMoE’s predicted cell-type proportion pattern is consistent between different slides (Fig. 10a). Furthermore, we show that scMoE predicts distinct cell-type proportions for non-malignant, DCIS, and IDC regions (Fig. 10b). scMoE predicts a high proportion of fibroblasts, as well as increased heterogeneity for IDC regions compared to DCIS regions. This is notable as IDC regions are noted to have elevated extracellular matrix associated expression [56]. Unlike the IDC spots, the DCIS spots are predicted to be comparatively homogeneous, with predominantly luminal and myoepithelial cells, also concordant with the established literature [57, 58].

As future work we will extend scMoE in several directions. The first would be to extend scMoE to cell-anchored single-cell multi-omics integration, such as scRNA+scATAC-seq or CITE-seq. The next challenge would be to extend the model to integrate multi-omics datasets that are not mutually cell-anchored, in either a mosaic integration setting or a more challenging diagonal integration setting with no overlapping features between datasets [59]. A current limitation of scMoE is that it doesn’t utilize the linear batch correction factors present in scETM. An additional future extension of scMoE would be to incorporate batch correction vectors, potentially by simultaneously learning batch correction factors in parallel across experts.

In summary, scMoE is a flexible and interpretable, hierarchical mixture of experts single-cell embedded topic model. By pre-training cell-type specific experts, we are able to improve downstream analysis and conduct CTS differential tests. We report quantitative results competitive with or surpassing existing models on 9 single-cell datasets and 3 simulated spatial bulk datasets. We additionally show biologically meaningful results on 3 bulk cancer datasets, a ST brain dataset, and a ST breast cancer dataset.

## 5 Acknowledgement

This work is supported by Canada Research Chair (Tier 2) in Machine Learning for Genomics and Healthcare (CRC-2021-00547), NSERC Discovery Grant (RGPIN-2016-05174), and New Frontier Research Fund – Exploration (NFRFE-2019–00980).

## S1 Additional Figures

**Figure S1:**
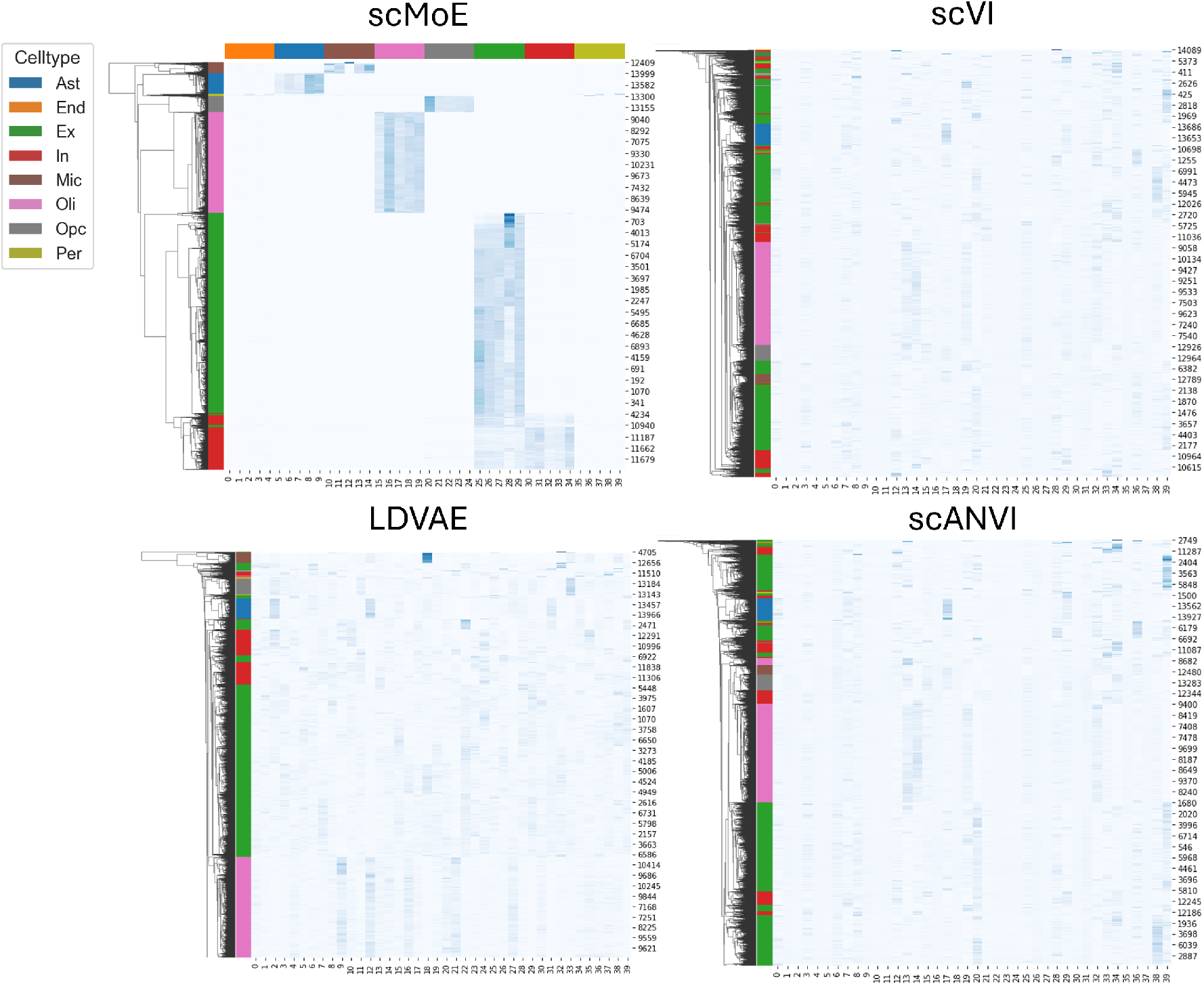
AD cross validation fold 1 left out cells normalized embedding heatmaps. Row colors correspond to cell-type labels. Column colors for scMoE panel indicate cell-type associated topics. scVI, LDVAE, and scANVI latent embeddings were normalized using Softmax to make visualizations comparable.

**Figure S2:**
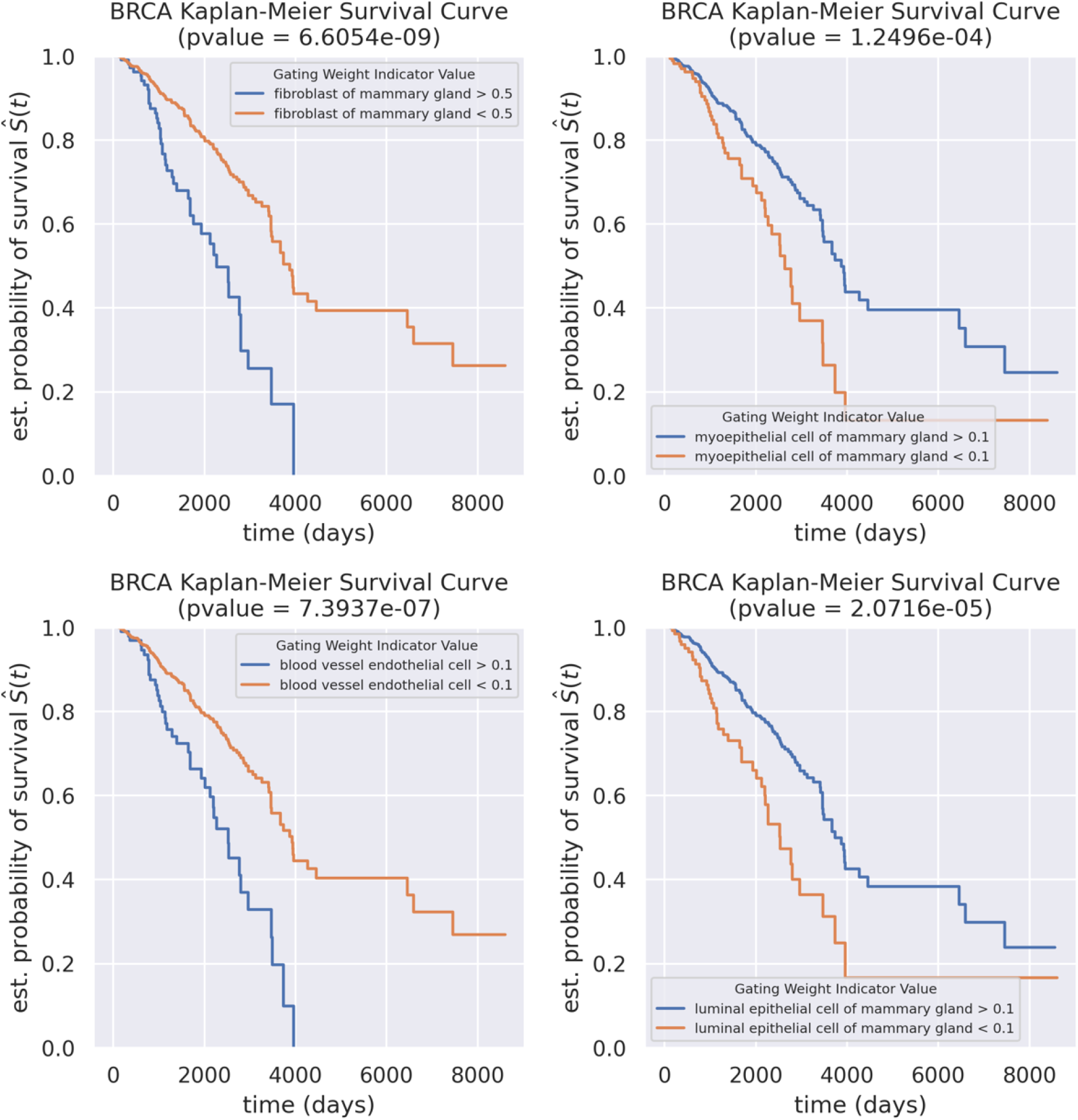
TCGA breast cancer survival curves, computed using thresholded scMoE predicted cell-type proportions. Cell-types of survival importance with multiple hypothesis testing adjusted p-values *<* 0.05 reported.

**Figure S3:**
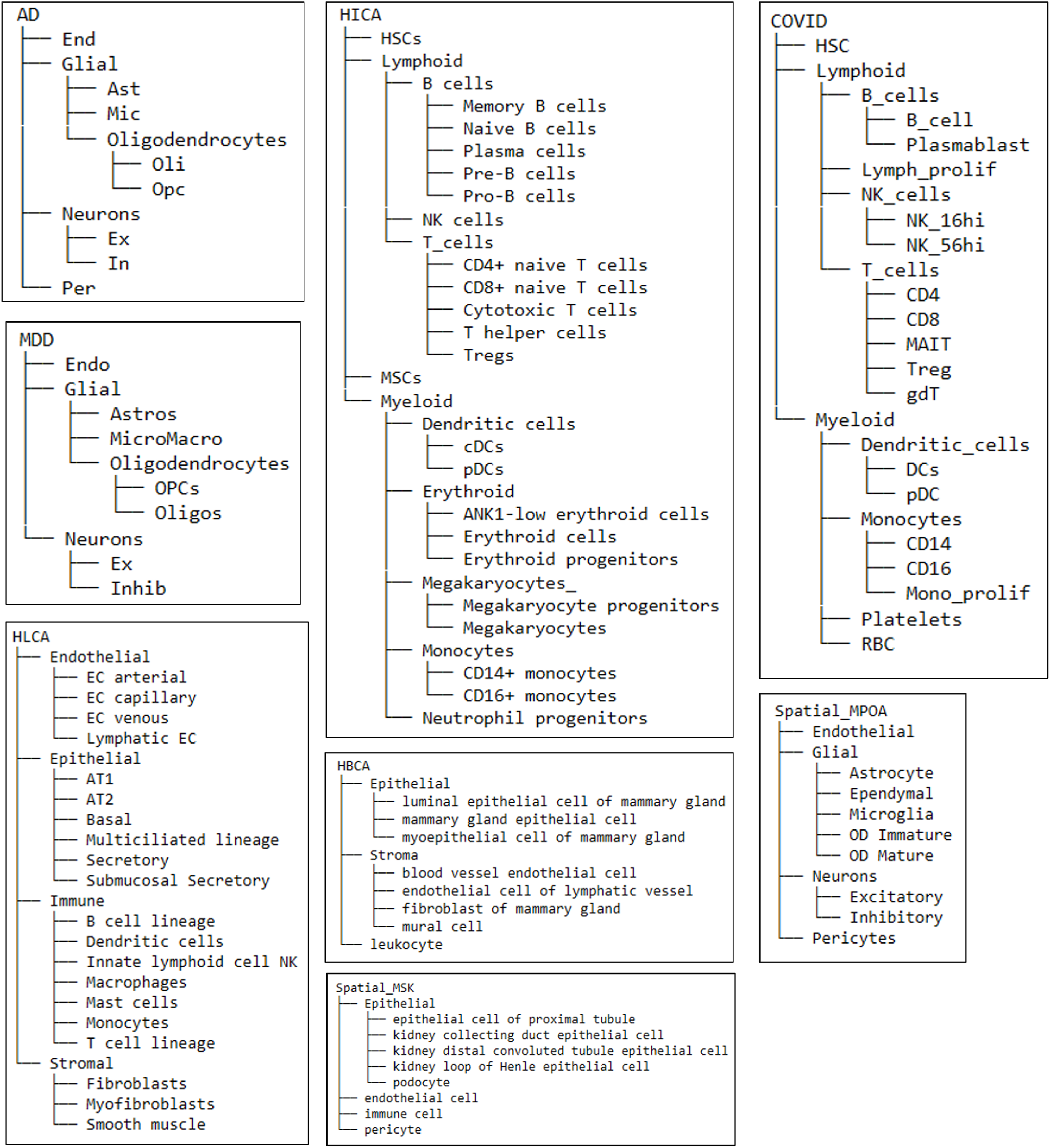
Hierarchical architectures for AD, MDD, HLCA, HICA, COVID, HBCA, MPOA and MSK.

**Figure S4:**
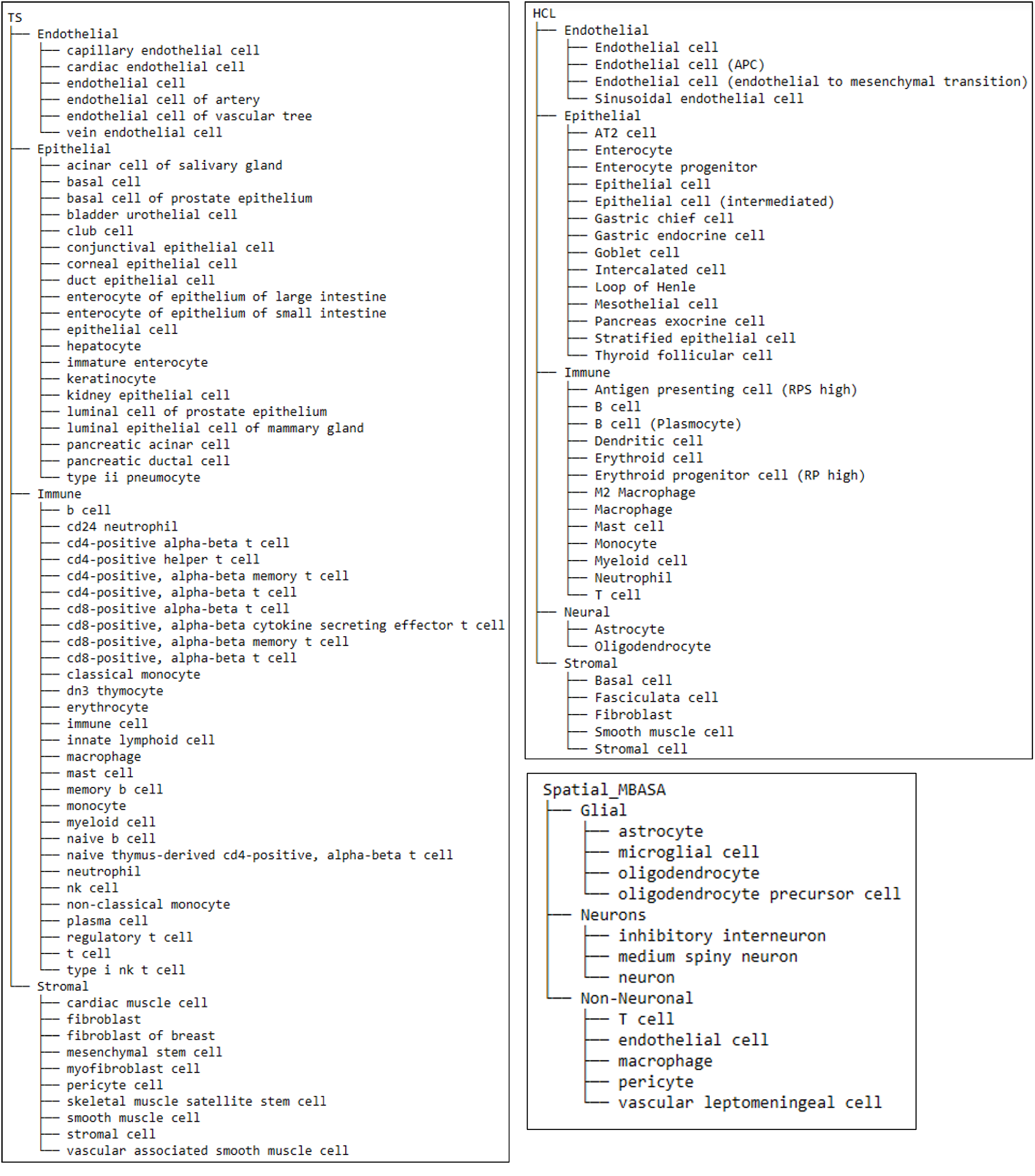
Hierarchical architectures for TS, HCL, and MBASA.

**Table S1:** Single-cell RNA-seq, bulk RNA-seq, and spatial transcriptomics dataset accession information and details.

| **Dataset** | **Data Type** | **Accession** | **# Cells / Samples** | **# Genes** |
| --- | --- | --- | --- | --- |
| Alzheimer's Disease | Single-cell | SYN18485175 | 70634 | 17775 |
| Adult Human Heart | Single-cell | ERP123138 | 451513 | 30321 |
| COVID19 Atlas | Single-cell | EMTAB10026 | 647366 | 24304 |
| Human Breast Cell Atlas | Single-cell | EMTAB13664 | 775610 | 33084 |
| Human Cell Landscape | Single-cell | GSE134355 | 330067 | 26942 |
| Hematopoietic Immune Cell Atlas | Single-cell | ERP122984 | 824385 | 24289 |
| Human Lung Cell Atlas | Single-cell | 10.5281/zenodo.7599104 | 583769 | 28021 |
| Major Depressive Disorder | Single-cell | GSE144136 | 73464 | 28788 |
| Tabula Sapiens | Single-cell | GSE201333 | 388382 | 4948 |
| TCGA Pancreatic Adenocarcinoma | Bulk | TCGA-PAAD | 181 | 18127 |
| TCGA Lung Adenocarcinoma | Bulk | TCGA-LUAD | 600 | 60660 |
| TCGA Acute Myeloid Leukemia | Bulk | TCGA-AML | 3225 | 60660 |
| TCGA Breast Cancer | Bulk | TCGA-BRCA | 1212 | 20502 |
| Dorsolateral Prefrontal Cortex | Spatial | jhpce#HumanPilot10x | 47681 (Spots) | 33538 |
| Spatial Breast Cancer | Spatial | 10.1126/SCIENCE.AAF2403 | 1031 (Spots) | 16385 |
| Mouse Medial Preoptic Area | MERFISH | 10.5061/dryad.8t8s248 | 64373 | 156 |
| Mouse Brain Aging Spatial Atlas | MERFISH | GSE207848 | 378708 | 374 |
| Mouse Spatial Kidney | MERFISH | 10.6084/m9.figshare.19310672.v2 | 126547 | 307 |

